# Decoding CRISPR–Cas9 PAM recognition with UniDesign

**DOI:** 10.1101/2023.01.08.523136

**Authors:** Xiaoqiang Huang, Jun Zhou, Dongshan Yang, Jifeng Zhang, Xiaofeng Xia, Y. Eugene Chen, Jie Xu

## Abstract

The critical first step in CRISPR–Cas mediated gene editing is recognizing a preferred protospacer adjacent motif (PAM) on target DNAs by the protein’s PAM-interacting amino acids (PIAAs). Thus, accurate computational modeling of PAM recognition is useful in assisting CRISPR–Cas engineering to relax or tighten PAM requirement for subsequence applications. Here we describe a universal computational protein design framework (UniDesign) for designing protein–nucleic acid interactions. As a proof of concept, we applied UniDesign to decode the PAM–PIAA interactions for eight Cas9 proteins. We show that, given native PIAAs, the UniDesign-predicted PAMs are largely identical to the natural PAMs of all Cas9s. In turn, given natural PAMs, the computationally redesigned PIAA residues largely recapitulated the native PIAAs (>70% and >80% in terms of identity and similarity, respectively). These results demonstrate that UniDesign faithfully captures the mutual preference between natural PAMs and native PIAAs, suggesting it as a useful tool for engineering CRISPR–Cas and other nucleic acid–interacting proteins. UniDesign is open-sourced at https://github.com/tommyhuangthu/UniDesign.

## Introduction

CRISPR–Cas mediated genome editing shows great promise in biotechnology and medicine (*1–5*). However, the specific recognition of protospacer adjacent motifs (PAMs) prior to DNA cleavage limits the target range of Cas nucleases (*6*). For instance, the most widely used SpCas9 from *S. pyogenes* recognizes an NGG (N is A/C/G/T) PAM, allowing it to target about 1/8 (¼ ×¼ × 2 DNA strands) human genomic sequences. Engineering the PAM requirement of a CRISPR–Cas protein has immediate applications (*7–10*). For example, relaxing the PAM requirement (e.g., from NGG to NGN) would increase the overall targetable sequences; whereas tightening the PAM requirement (e.g., from NGG to AGG) would increase editing specificity and reduce off-target edits (*6*).

Directed evolution and structure-guided engineering are the two prevalent strategies to engineer the PAM preference of CRISPR–Cas proteins. In recent years, the latter has gained more popularity and success along with the increased number of solved Cas protein structures. Most of these efforts, however, are limited to utilizing the structural biology knowledge to identify key PAM-interacting amino acids (PIAAs) for mutagenesis engineering.

By contrast, computational protein design (CPD) approaches make use of protein structures as a basis to interrogate the effects of mutagenesis with advanced computational algorithms (*11–13*) and at a higher level to de novo design a protein sequence with desired function (*14, 15*). CPD methods have been widely used in protein engineering, and yielded proteins with improved functional characteristics (*12, 16–18*). However, the powerful CPD methods were rarely adapted to engineer Cas9 protein to modify their PAM requirements. The COMET workflow is by far the only computational approach to model PAM recognition (*19*). COMET provided a computational interpretation to the KKH variant of *S. aureus* Cas9 (SaCas9), which was obtained by directed evolution that relaxes the PAM from NNGRRT into NNNRRT (*10*), followed by COMET guided design of two new variants SaCas9-NR and SaCas9-RL, both with a further relaxed PAM of NNGRRN (*19*). COMET leveraged molecular dynamics (MD) simulation and free-energy perturbation (FEP) to calculate mutation-induced binding free energy change, which was subsequently correlated with PAM recognition (*19*). One caveat of COMET is that both MD and FEP are computationally expensive and resource-inefficient, limiting its large-scale application and adaptation.

We previously developed two CPD methods, namely EvoDesign and EvoEF2, for monomer protein design and protein–protein interaction (PPI) design (*20, 21*). EvoDesign is an evolution-based approach that combines evolutionary profile (represented as position-specific scoring matrix, PSSM) derived from multiple structure/sequence alignment (MSSA) with a physics-based energy function (EvoEF) for protein sequence design (*20*). In some cases, the PSSM term may not be reliable due to insufficient structure analogs obtained, while the accuracy of EvoEF by itself is moderate for de novo protein design (*21*). To increase the physical energy function’s accuracy, we developed EvoEF2 by introducing a few statistical energy terms into EvoEF and reoptimized all the energy weights through extensive de novo sequence design (*21*). EvoEF is a linear combination of five terms, including van der Waals, hydrogen bonding, electrostatics, solvation, and reference energy, and optimized by maximizing the accuracy of predicting the thermodynamic changes upon mutations (*20*). Compared with EvoEF, EvoEF2 includes four extra knowledge-based terms, i.e., disulfide-bonding, amino acid propensity, Ramachandran, and rotamer frequency. Benchmark results showed that all nine terms are important to EvoEF2’s high accuracy, and overall, EvoEF2 improved the sequence design accuracy by about two folds relative to EvoEF (*21*). We have successfully used these two methods to design novel protein and peptide binders (*22, 23*).

Unlike COMET which relies on resource-heavy algorithms (i.e., MD and FEP), both EvoDesign and EvoEF2 methods comprise resource-friendly components: a rotamer library for efficient amino-acid conformation sampling, an efficient energy function for protein sequence scoring, and a fast optimization algorithm for searching low-energy designer sequences (*21, 24, 25*). With more approximations, these two methods are much faster than MD-based approaches yet still very accurate, making them particularly appropriate to efficiently explore the vast sequence space, such as those of the CRISPR–Cas proteins.

Here we report the development of a universal CPD framework (UniDesign) that is built on EvoDesign and EvoEF2 by adding a new capacity to model and design the protein–nucleic acid interaction (PNI), a functionality that is necessary yet underdeveloped for engineering CRISPR–Cas protein’s PAM preference. Our detailed computational analyses show that UniDesign captures the “self-consistency” between PIAAs and PAMs satisfactorily, and provides computational explanation at the molecular and energetic levels, thus suggest the potential application of UniDesign in assisting CRISPR–Cas protein engineering.

## Methods

### Cas9 structures collection and preprocessing

The PDB files for eight Cas9 proteins were downloaded from RSCB PDB (*26*). Ions and water molecules were removed except for 5AXW, 5CZZ, and 6JDV, in which PAM-interacting water molecules were retained. Specifically, HOH1202, HOH1217, HOH1227, HOH201, HOH202, and HOH203 were kept in 5AXW. HOH1207, HOH1233, HOH201, HOH202, HOH203, and HOH204 were reserved in 5CZZ. HOH1201 was retained in 6JDV. The missing amino-acid side chains were added using the “RepairStructure” command in UniDesign, and sidechain steric clashes were reduced using the UniDesign “Minimization” command.

### Atomic parameters and topologies for nucleotides

The amino-acid atomic parameters and topologies in EvoEF2 (and UniEF) are adapted from the united-atom force field CHARMM19 (*21*). However, there is no parameters and topologies for nucleic acids in CHARMM19. We took nucleotide topologies from the all-atom CHARMM36 force field (*28*) but removed all nonpolar hydrogen atoms. To be consistent with nucleotide nomenclature in PDB, the nucleotides were renamed as DA, DC, DG, and DT for DNA, and A, C, G, and T for RNA in the UniEF energy function. The nucleotide atoms were parameterized based on their similarity to atoms in amino acid chemical groups. Our previous study demonstrated the good performance of CHARMM19-based EvoEF2 on protein sequence design (*27*), rationalizing the development of UniEF using CHARMM19-like parameters and topologies. The UniEF atomic parameters and residue topologies can be found at https://github.com/tommyhuangthu/UniDesign/blob/master/library/toppar/param_charmm19_lk.prm and https://github.com/tommyhuangthu/UniDesign/blob/master/library/toppar/top_polh19.inp, respectively.

### UniEF and modification to the hydrogen-bonding energy term

In general, UniEF inherits the EvoEF2 energy function for protein design (*21*). UniEF is the linear combination of nine energy terms:

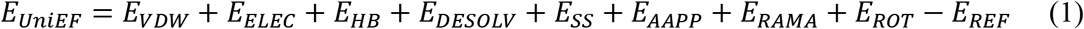

Here, *E_VDW_*, *E_ELEC_, E_HB_, E_DESOLV_*, and *E_SS_* represents the total van der Waals, electrostatic, hydrogen-bonding, de-solvation, and disulfide-bonding interaction, respectively; these terms are calculated as the weighted sum of pairwise atomic interaction energy. *E_AAPP_, E_RAMA_*, and *E_ROT_* represents the term for calculating amino acid propensity, the Ramachandran term, and the term for modeling rotamer frequency in the rotamer library, respectively; these terms are dependent on protein backbone geometry, and are calculated as the sum of residue- or rotamer-wise energy. Finally, *E_REF_*, namely protein reference energy, models the energy of a protein in the unfolded state, and is roughly calculated as the sum of amino acid reference energy.

Compared to EvoEF2, we updated the hydrogen-bonding energy term in UniEF to model water-mediated hydrogen bonds, which are commonly seen in biological systems. For a regular hydrogen-bonding interaction, *E_HB_*(*D,H,A,B*) is defined by four atoms: the hydrogen atom (*H*), the hydrogen acceptor (*A*), the hydrogen donor (*D*), and the base atom to which *A* is attached (*B*). *E_HB_*(*D,H,A,B*) is a linear combination of three terms that depend on the distance between *H* and *A* (*d_HA_*), the angle between *D*, *H*, and *A* (*θ_DHA_*), and the angle between *H*, *A*, and *B* (*φ_HAB_*):

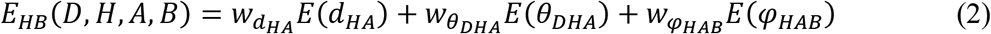

where:

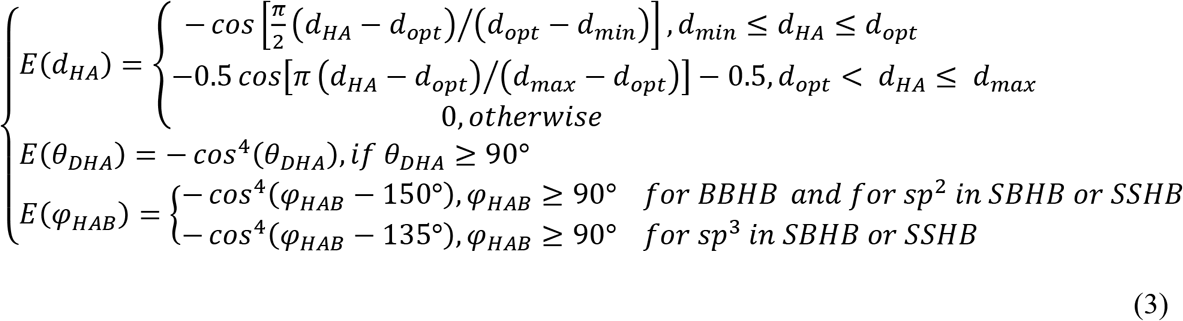

*w_d_HA__, w_θ_DHA__*, and *w_φ_HAB__* are weights for the three terms. The optimal distance between *H* and *A, d_opt_*, is set to 1.9 Å. Additionally, *d_min_* = 1.4 Å and *d_max_* = 3.0 Å are the lower and upper bounds of the distance between the hydrogen–acceptor pair. The optimal *φ_HAB_* value is set to either 150° or 135°, depending on the acceptor hybridization (sp^2^ or sp^3^) and the locations of the donor and acceptor atoms (BBHB: Backbone-Backbone Hydrogen Bond; SBHB: Sidechain-Backbone Hydrogen Bond; SSHB: Sidechain-Sidechain Hydrogen Bond).

Water molecules can be a hydrogen bond donor, acceptor, or both. In UniDesign, the water hydrogen atoms are not explicitly modeled due to the difficulty to determine their positions. Water-mediated hydrogen bonding interactions are treated in three cases:

Case 1: the hydrogen bond acceptor (*A*) is water while the donor is normal. In this case, the base atom *B* does not exist, and hence *φ_HAB_* related terms in Equations 2 and 3 are omitted. The other terms are the same.
Case 2: the hydrogen bond donor (*D*) is water while the acceptor is normal. In this case, the hydrogen atom (*H*) is omitted, and *E_HB_*(*D,A,B*) is used to calculate water-mediated hydrogenbonding energy.

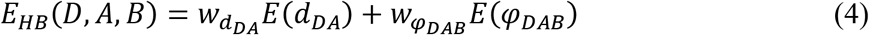

where:

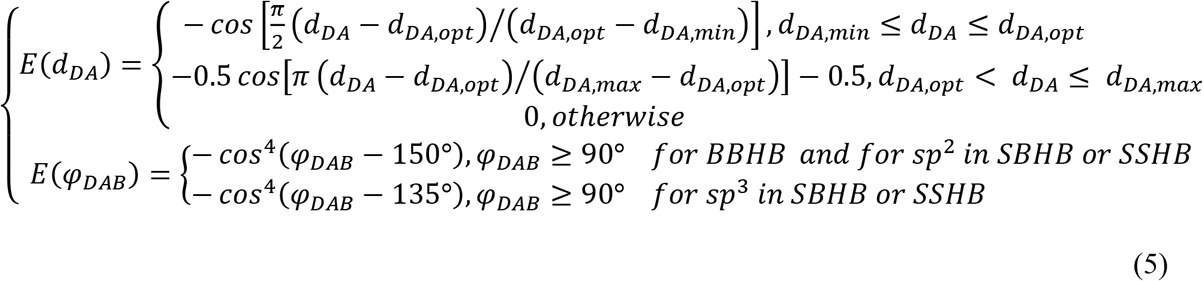 The optimal distance between *D* and *A, d_DA,opt_*, is set to 2.8 Å. The minimum and maximum distances for considering a water-mediated hydrogen bond are: *d_DA,min_* = 2.3 Å and *d_DA,max_* = 3.9 Å.
Case 3: both *D* and *A* are water. In this case, *H* and *A* do not exist, and only the *E*(*d_DA_*) term in Equations 4 and 5 is used to calculate water-mediated hydrogen bonding energy.

### Generation of PAM combo models with UniDesign

The “BuildMutant” command, implemented in EvoEF2 to build amino-acid mutant models, was extended to build models for nucleotide mutations in UniDesign. When mutating a nucleotide, the torsional angle centered on the bond that connects the backbone and sidechain is kept unchanged, and the coordinates of sidechain atoms are calculated based on the nucleotide topology described above. For instance, for the DA→DC mutation, the value of torsional angle will be used for the value of torsional angle O4’–C1’–N1–C2 in nucleotide DC will be taken from that of O4’–C1’–N9–C4 in nucleotide DA. For paired DNA strands, if one nucleotide is mutated on one strand, the paired nucleotide on the other strand will be automatically mutated to ensure reverse complementarity. Below is the command line to build PAM mutants using 4UN3 as an example:

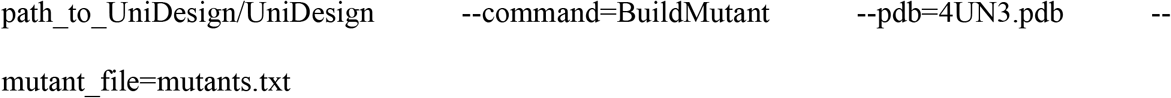

The “*mutants.txt*” file contains one or more lines like “*tD5a,gD6a,gD7a*;”. Each line, ended with “;”, represents one mutant for which mutations are separated by “,”. In this example, the TGG PAM will be mutated into AAA. All other PAM combos can be built similarly.

### Computational repacking and redesign of PIAAs

The UniDesign “ProteinDesign” command was used to repack or redesign the PIAA residues. Below is the command line using 4UN3 as an example:

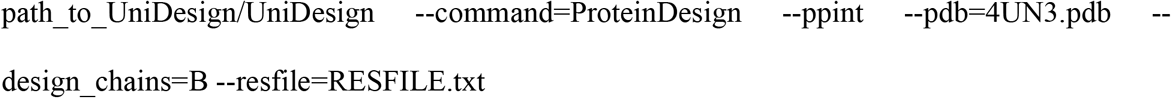

The “--*ppint*” option specifies a PPI or PNI design task which is equally treated in the prototype UniDesign. The “--*design_chains=B*” option means that design is carried out on chain B, i.e., the SpCas9 protein chain. The residues to be repacked and/or redesigned were controlled using a restraint file (RESFILE) “*RESFILE.txt*”. The RESFILE format is explained in detail at https://github.com/tommyhuangthu/UniDesign/blob/master/manual.docx.

## Results

### UniDesign for protein–nucleic acid interaction modeling and design

EvoDesign and EvoEF2 lack the capacity for PNI design or other functional protein design tasks like protein–ligand interaction (PLI) design and enzyme design, since they cannot model non-protein molecules. To overcome these limitations, we have developed a universal CPD approach named UniDesign to deal with these four kinds of functional protein design tasks (i.e., PPI, PNI, PLI, and enzyme design) (**Fig. 1a**).

**Fig. 1.**
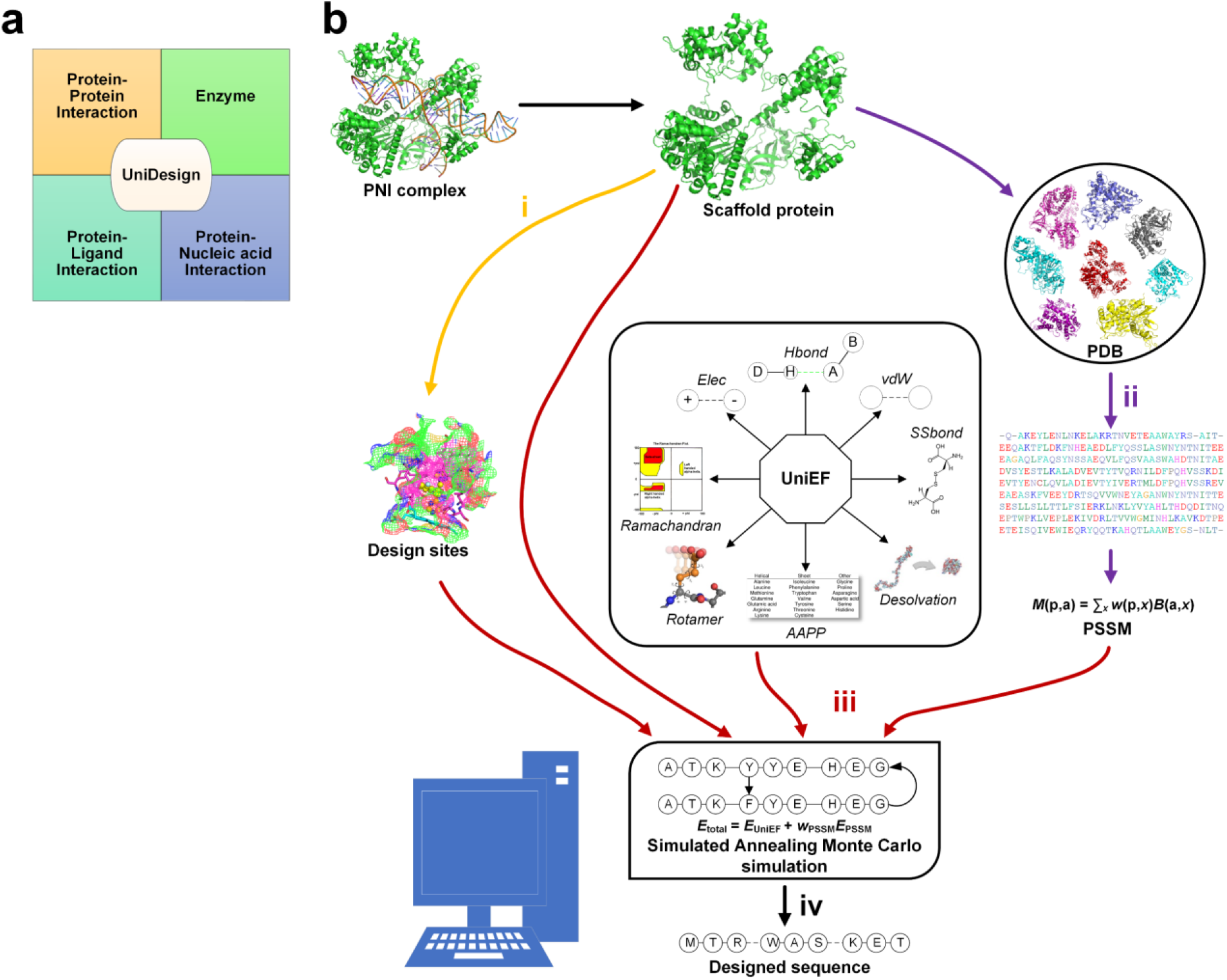
The UniDesign workflow for protein–nucleic acid interaction design. (**a**) UniDesign can work for protein–protein interaction design, protein–ligand interaction design, enzyme design, and protein–nucleic acid interaction (PNI) design. (**b**) The UniDesign pipeline for PNI design comprises of four stages (denoted as i, ii, iii, and iv).

UniDesign inherits the overall methodology of EvoEF2 while adopting the evolutionary component of EvoDesign. **Fig. 1b** illustrates the pipeline for PNI design. First, design sites of interest can be defined based on the input PNI complex structure. Second, UniDesign searches for structure analogs to the scaffold protein and constructs a PSSM using the MSSA obtained from the pairwise structure alignment. Third, building on the PNI scaffold, UniDesign performs sequence redesign for the predefined design sites, using a composite energy function by combining the *E_PSSM_* and *E_UniEF_* energy terms, where UniEF is a pretrained energy function extended from EvoEF2 (*21*), and is used to model the physical interactions between protein and nucleic acid (see Methods). An efficient simulated annealing Monte Carlo (SAMC) simulation procedure is employed to search the sequence space. Finally, after SAMC simulation, the sequence with the lowest total energy is taken as the best design. Due to SAMC’s stochasticity, the lowest-energy designs from multiple independent simulations will be collected for analysis. In UniDesign, the evolutionary module is set as optional; UniEF alone is used for design when evolution is disabled.

### Summary of eight CRISPR–Cas9 proteins in this study

We selected eight CRISPR–Cas9 proteins, namely SpCas9, SaCas9, FnCas9, Nme1Cas9, Nme2Cas9, CdCas9, St1Cas9, and AceCas9 (**Table 1**), to study their PAM recognition because (1) their experimentally determined consensus PAMs are consistent in different studies, providing a ground truth to computational modeling; (2) their Cas9/gRNA (guide RNA)/DNA complex structures are available with PDB (Protein Data Bank) PAM being a part of the consensus PAMs (**Table 1**) [PAM sequence is 5’ to 3’ on non-target strand (NTS)]. The “consensus” PAMs refer to the preferred PAMs that are associated with high cleavage efficiency.

**Table 1.** Summary of the PAMs for eight Cas9 proteins.

| Cas9 | Species | Length | Consensus PAM | PDB (resolution) | PDB PAM |
| --- | --- | --- | --- | --- | --- |
| SpCas9 | *S. pyogenes* | 1,368 | NGG | 4UN3 (2.59 Å) | TGG |
|  |  |  |  | 5F9R (3.40 Å) | TGG |
| SaCas9 | *S. aureus* | 1,053 | NNGRRT | 5AXW (2.70 Å) | TTGGGT |
|  |  |  |  | 5CZZ (2.60 Å) | TTGAAT |
| FnCas9 | *F. novicida* | 1,629 | NGG | 5B2O (1.70 Å) | TGG |
| Nme1Cas9 | *N. meningitidis* | 1,082 | NNNNGATT | 6JDV (3.10 Å) | ATATGATT |
| Nme2Cas9 | *N. meningitidis* | 1,082 | NNNNCC | 6JE3 (2.93 Å) | AGGCCC |
| CdCas9 | *C. diphtheriae* | 1,084 | NNRHHHY | 6JOO (2.90 Å) | GGGTAAT |
| St1Cas9 | *S. thermophilus* | 1,121 | NNRGAA | 6M0W (2.76 Å) | AAAGAA |
| AceCas9 | *A. cellulolyticus* | 1,138 | NNNCC | 6WBR (2.91 Å) | ATACC |

For SpCas9, two structures determined at different resolutions, which represent the catalytically inactive and active state respectively, were chosen (PDB ID: 4UN3 and 5F9R). For SaCas9, two structures (5AXW and 5CZZ) with different PDB PAMs were considered. One structure was used for each of the remaining six Cas9 proteins (i.e., FnCas9, Nme1Cas9, Nme2Cas9, CdCas9, St1Cas9, and AceCas9). We did not consider the minimal Cas9 from *C. jejuni* (CjCas9, 984 amino acids) because its PAM determined in different studies are not consistent (*29–32*).

Protein structures provide direct insights to understand how Cas9 amino acids recognize PAMs. In the two structures of SpCas9 (*33, 34*), whose PAM preference is NGG, it is noted that while the second G and third G of the PDB PAM form bidentate hydrogen bonds with Arg1333 and Arg1335 respectively, the first T does not form direct contacts with amino acids (**Fig. 2a** and **b**).

**Fig. 2.**
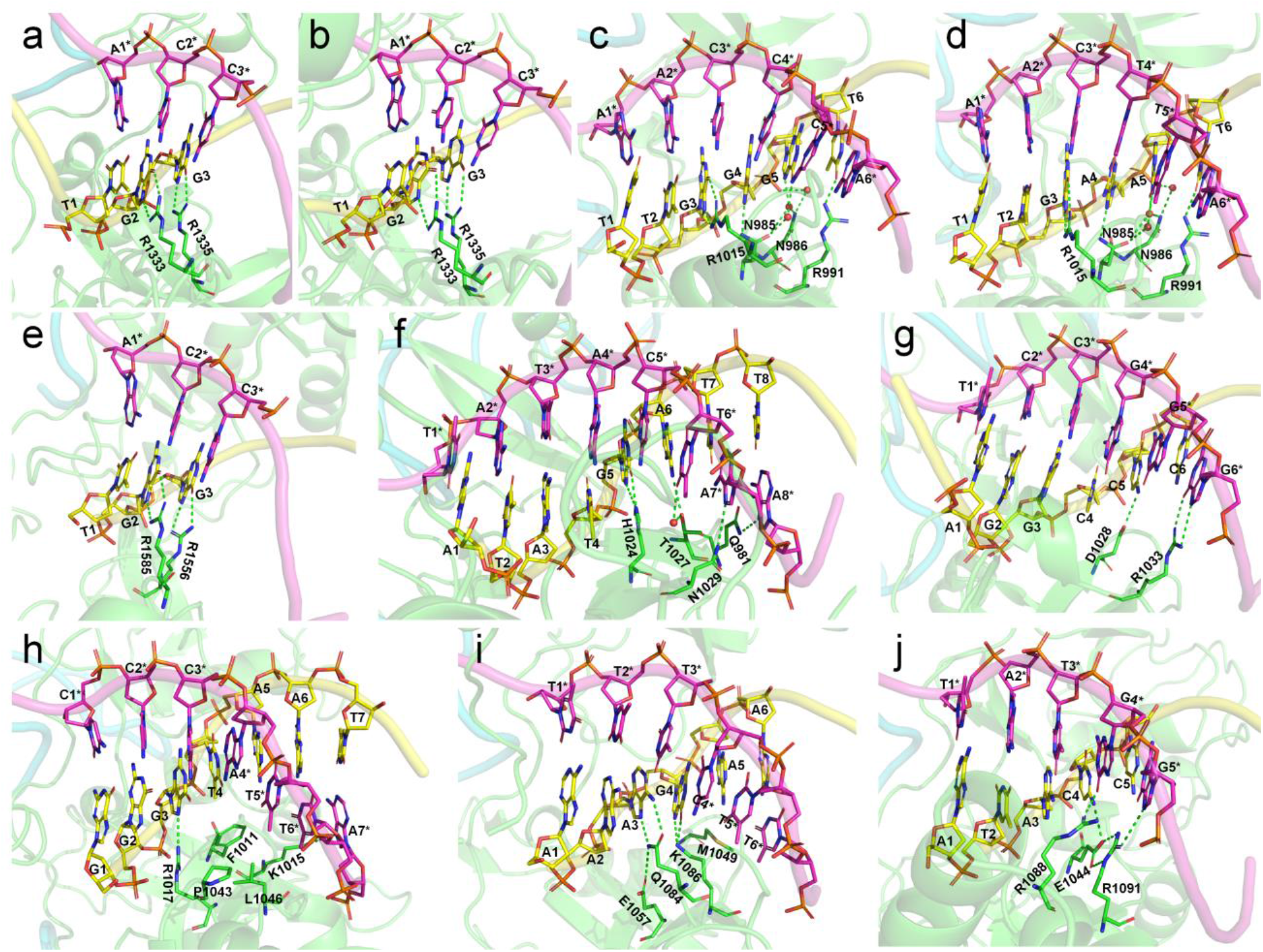
Visualization of the interactions between PAM nucleotides and PAM-recognizing amino acids. (**a**) 4UN3 (SpCas9) with TGG PAM; (**b**) 5F9R (SpCas9) with TGG PAM; (**c**) 5AXW (SaCas9) with TTGGGT PAM; (**d**) 5CZZ (SaCas9) with TTGAAT PAM; (**e**) 5B2O (FnCas9) with TGG PAM; (**f**) 6JDV (Nme1Cas9) with ATATGATT PAM; (**g**) 6JE3 (Nme2Cas9) with AGGCCC PAM; (**h**) 6JOO (CdCas9) with GGGTAAT PAM; (**i**) 6M0W (St1Cas9) with AAAGAA PAM; (**j**) 6WBR (AceCas9) with ATACC PAM. Nucleotides on nontarget and target strands are shown as yellow and magenta sticks respectively; nontarget-stranded nucleotides are marked with asterisks. Amino acids are shown in green sticks. Hydrogen bonds are shown as green dashed lines.

Similarly, in the two structures of SaCas9 (*35*), which has a PAM preference of NNGRRT, the first two T’s of the PDB PAM do not form direct interactions or water-mediated interactions with amino acids; whereas the third G forms bidentate hydrogen bonds with Arg1015, the fourth G or A forms one hydrogen bond with Asn985, the fifth G or A forms two water-mediated hydrogen bonds with Asn985 and Asn986 respectively, and the sixth T forms a hydrogen bond with Arg991. Besides, Asn986 forms a water-mediated hydrogen bond with the phosphate oxygen atoms of the fifth G or A (**Fig. 2c** and **d**).

FnCas9 was reported to recognize both NGG and NGA with a much higher preference of NGG (*36*). In the structure of FnCas9, the second G forms coplanar bidentate hydrogen bonds with Arg1585. The third G or A can respectively form distorted bidentate hydrogen bonds or a single hydrogen bond with Arg1556 (**Fig. 2e**).

Nme1Cas9 and Nme2Cas9 are type-II Cas9 proteins in which both DNA strands are involved in PAM recognition (*37*). Nme1Cas9 was reported to recognize promiscuous PAMs with a consensus on NNNNGATT (*38–40*). Nme2Cas9 has a PAM preference for NNNNCC (*41*). In the structure of Nme1Cas9 (*37*), the fifth G forms two hydrogen bonds with His1024. The sixth A forms one hydrogen bond with Thr1027, while its paired T can form a water-mediated hydrogen bond with Thr1027. The seventh and eighth T’s of PAM do not have contacts with amino acids, but the paired A’s on the target strand (TS) form one and two hydrogen bonds with Asn1029 and Gln981, respectively (**Fig. 2f**). In the structure of Nme2Cas9 (*37*), the fifth C forms a hydrogen bond with Asp1028. The sixth C does not form contacts with amino acids but the paired G forms bidentate hydrogen bonds with Arg1033 (**Fig. 2g**).

CdCas9 is also a type-II Cas9 protein with a PAM preference for NNRHHHY (*42*). In its reported structure (*42*), the third G forms a hydrogen bond with Arg1017. The fourth to seventh PAM nucleotides (TAAT) do not form any hydrogen bond with amino acids. The fourth T and the fifth–sixth A’s paired partners (two T’s) form van der Waals contacts with Phe1011, Pro1043, Leu1046, and Lys1015. The seventh T’s paired A forms a hydrogen bond with Lys1015 (**Fig. 2h**).

In the structure of St1Cas9 (*43*), whose PAM preference is NNRGAA, the third A forms bidentate hydrogen bonds with Gln1084 which is positioned by Glu1057. The fourth G forms two hydrogen bonds with Lys1086. The fifth A and the T in pair with the sixth A form van der Waals interactions with Met1049 (**Fig. 2i**).

In the structure of AceCas9 (*44*), whose PAM preference is NNNCC, the fourth C forms a hydrogen bond with Glu1044, which is positioned by Arg1091, and the paired G forms a hydrogen bond with Arg1088. The fifth C’s paired G on the TS forms bidentate hydrogen bonds with Arg1091 (**Fig. 2j**).

In sum, these reported CRISPR–Cas9 structures provide direct visual clues to the key interactions, commonly through hydrogen bonds or van der Waals forces, between PIAAs and PAMs.

### Computational modeling reveals that native PIAAs prefer consensus PAMs

We proceeded to develop a CPD method to study the interaction between the CRISPR–Cas protein’s PIAAs and their preferred PAMs on the target DNAs. We defined the amino acids that are in direct contacts (<4.5 Å) with the side chains of PAMs or the paired nucleotides as PIAAs (**Supplementary Table 1**). Of note, PIAAs defined by this simple rule contained all PAM-recognizing residues described in the above section.

For each PDB, its PAM nucleotides were mutated to generate all *4^L^* PAM combos, where *L* is PAM length. For example, the consensus PAM of SpCas9 is NGG, and we generated all 64 (= 4^3^) combos from AAA to TTT. The number of combos increases exponentially as *L* grows. Too many structures need to be generated for very long PAMs, e.g., 16,384 and 65,536 structures for CdCas9 and Nme1Cas9 that have seven and eight PAM nucleotides, respectively. Thus, only the last six PAM positions were varied if *L* > 6, resulting in a handleable maximum of 4,096 structures (this is reasonable because the first two nucleotides form no contacts with amino acids in CdCas9 or Nme1Cas9). When PAM nucleotides on NTS were mutated, the paired nucleotides on TS were correspondingly substituted to ensure complementarity.

We first computationally examined the preference of native Cas9 proteins to all PAM combos. This is to validate whether UniDesign can recapitulate such PAM preference for each of the eight Cas9 proteins. To do this, we used UniDesign to repack the PIAAs in the Cas9/gRNA/DNA complex model of each combo and calculate the total and binding energy. The “total energy” represents the energy of entire system including protein, gRNA, DNA, and water molecules (if applicable). The “binding energy” refers to the interaction between protein/gRNA and DNA, which is important to PAM recognition.

We reasoned that the binding energy is a good indicator for quantifying to what extent Cas9 prefers a PAM (a lower binding energy indicates a higher preference). Interestingly, for each Cas9, the energy associated with combos with consensus PAMs distributed in a relatively narrow window to the minimum binding energy 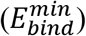 and/or the minimum total energy 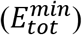 (**Fig. 3**). Through structure inspection, we noticed that some combos were scored a low binding energy at the cost of a high total energy (LbHt) because of inter-residue steric clashes. Mathematically, we defined a PAM as LbHt if its associated total energy is greater than a threshold (*δE_tot_*) of the minimum total energy 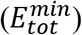. Because the PNI interactions between these LbHt PAMs and the corresponding Cas9s may not be physically stable/feasible, we filtered them out for subsequent computations.

**Fig. 3.**
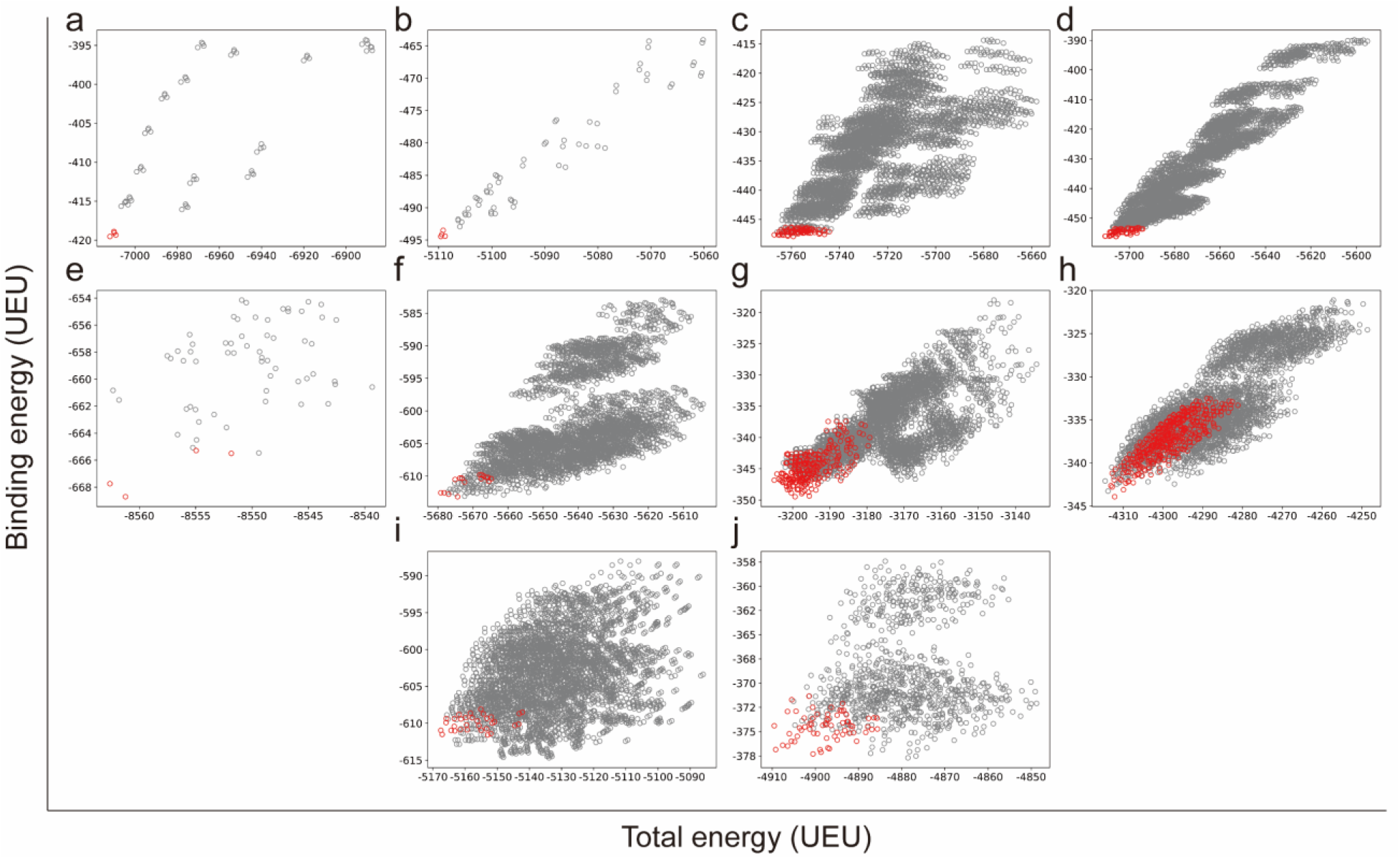
Total and binding energy for all PAM combos on different Cas9 scaffolds. (**a**) 4UN3 (SpCas9); (**b**) 5F9R (SpCas9); (**c**) 5AXW (SaCas9); (**d**) 5CZZ (SaCas9); (**e**) 5B2O (FnCas9); (**f**) 6JDV (Nme1Cas9); (**g**) 6JE3 (Nme2Cas9); (**h**) 6JOO (CdCas9); (**i**) 6M0W (St1Cas9); and (**j**) 6WBR (AceCas9). The combos with consensus PAMs are colored in red while the others in gray. UEU, UniDesign energy unit.

Only PAM combos that satisfy 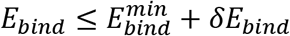 and 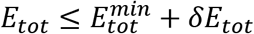 were subjected to sequence logo analysis using WebLogo (*45*) with appropriate *δE_bind_* and *δE_tot_* parameters; 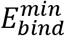 and 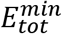 were yielded from UniDesign calculations (**Supplementary Table 2**). For comparison, sequence logos were also plotted with 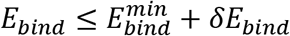 or 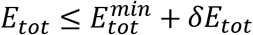 (**Supplementary Figs. 1** and **2**).

For both SpCas9 structures, the computationally determined preferred PAMs are NGG (**Fig. 4a** and **b**), consistent with the experimentally determined NGG consensus.

**Fig. 4.**
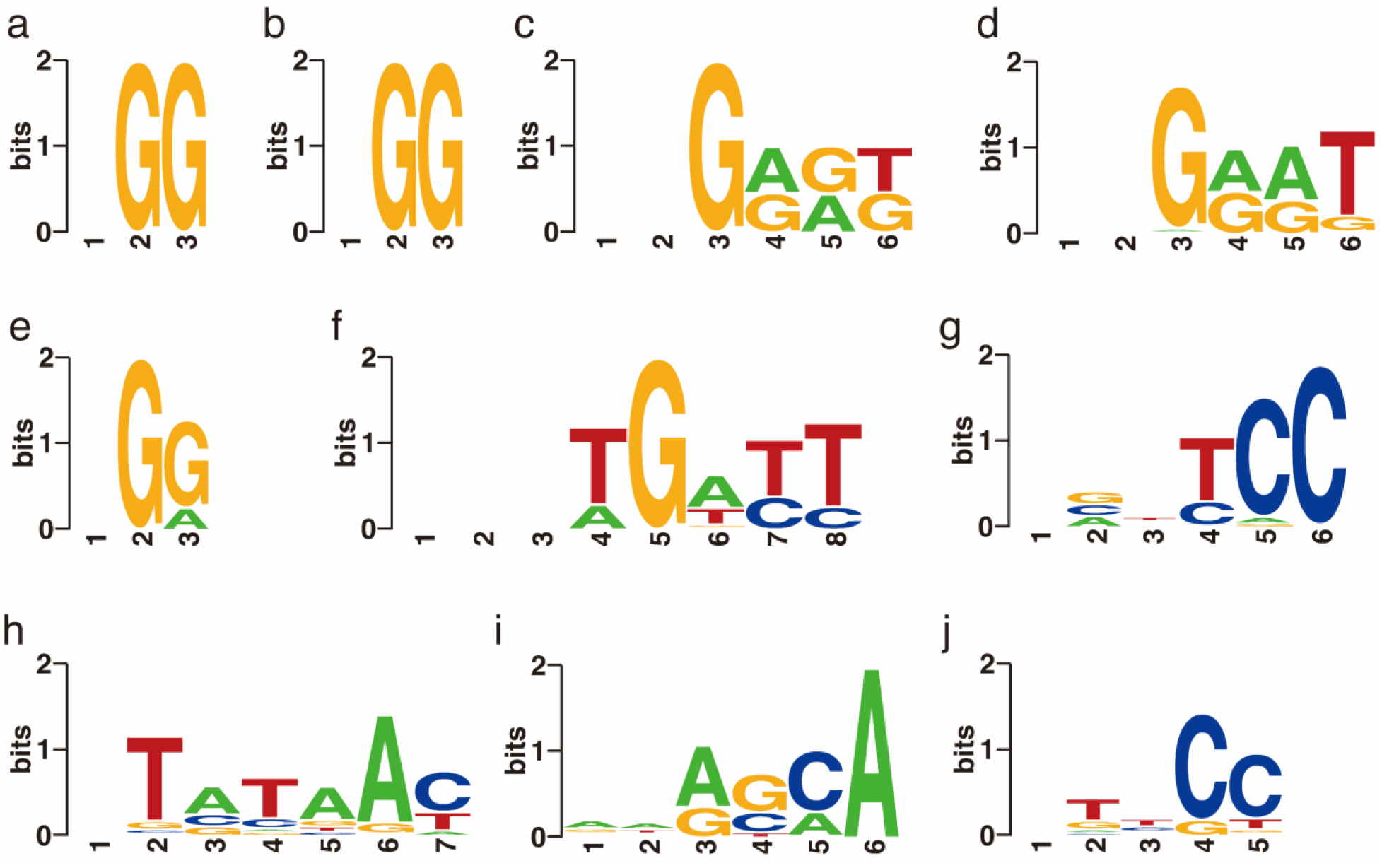
Computational modeling reveals that native Cas9 proteins prefer consensus PAMs. (**a**) result on scaffold 4UN3 (SpCas9); (**b**) result on scaffold 5F9R (SpCas9); (**c**) result on scaffold 5AXW (SaCas9); (**d**) result on scaffold 5CZZ (SaCas9); (**e**) result on scaffold 5B2O (FnCas9); (**f**) result on scaffold 6JDV (Nme1Cas9); (**g**) result on scaffold 6JE3 (Nme2Cas9); (**h**) result on scaffold 6JOO (CdCas9); (**i**) result on scaffold 6M0W (St1Cas9); and (**j**) result on scaffold 6WBR (AceCas9). Note that for Nme1Cas9 and CdCas9, the first two and one positions were excluded from PAM combo generation respectively, and the symbols were manually added at these positions for plotting sequence logos.

For both SaCas9 structures, the computationally determined PAM preference is NNGRR[T>G] (**Fig. 4c** and **d**), also matching the experimental consensus NNGRRT and the report that the sixth position tolerates other nucleotides albeit it prefers T most (*46, 47*).

For FnCas9, UniDesign obtains a computational preference for the NG[G>A] PAM (**Fig. 4e**), consistent with the experimentally determined PAM NG[G>A] (*36*).

For Nme1Cas9, the UniDesign computed its preferred PAM is NNNTGATT (**Fig. 4f**), which is a subset of the experimentally determined consensus NNNNGATT (*38–40, 48*). UniDesign modeling computationally confirmed that Nme1Cas9 prefers only G at the fifth position. Further, UniDesign predicted T, C, and C as the second favorable nucleotides at the sixth to eighth positions respectively. Besides, UniDesign suggested NNNNGACT, NNNNGCTT, NNNNGTCT, NNNNGTTT as alternative PAMs for Nem1Cas9 (**Fig. 4f**), which corroborate with the findings by Amrani et al. (*38*). We note that there is a discrepancy at the fourth PAM position where UniDesign favors T and A but the experimental studies do not indicate any preference. Interestingly, when the restraint on total energy was removed (e.g., *δE_tot_* = 1000) in UniDesign, the preference at the fourth PAM position is gone and the modeled consensus PAM becomes NNNNGNNT (**Supplementary Fig. 1**), the same as reported by Esvelt et al. in their study (*48*) (see Table 1 in the ref.).

For Nme2Cas9, the UniDesign-modeled PAM is N[G/C/A]N[T/C]CC with dominating C’s at the fifth and sixth positions (**Fig. 4g**), largely consistent with the experimental PAM NNNNCC (*37, 41*). UniDesign modeling suggests that the second position prefers equally G, C, and A but not T, in part due to the steric clashes between the 5-methyl group on T and Lys1044. Besides, computational modeling suggests that the fourth position prefers T or C probably because the paired A or G on the TS can form a potential hydrogen bond with Tyr1035.

For CdCas9, the UniDesign-modeled PAM is NTATAAY (**Fig. 4h**), a subset of the experimental consensus NNRHHHY (*42*). Computational modeling indicates that the third position prefers A > C ≈ G but not T, compared to the preference for G ≈ A > C but not T determined experimentally (*42*). The most discrepancy takes place at the second PAM position, where computation suggests a preference of T > G while experimental assay shows that all nucleotides are acceptable with a slight preference of G over others (*42*).

For St1Cas9, the UniDesign computed PAM is NNR[G ≈ C][C ≈ A]A (**Fig. 4i**), covering the experimentally determined consensus NNRGAA (*43*). In the computational model with a UniDesign suggested PAM ATACCA, the fourth C- and the fifth C-paired G’s formed hydrogen bonds with Ser1082 and K1086, respectively. Of note, the conformations of Ser1082 and K1086 in our UniDesign modeling by using the ATACCA PAM were quite different from those in the reported PDB model with a PAM of AAAGAA.

For AceCas9, UniDesign predicted the PAM preference as NTNCC (**Fig. 4j**), similar to the experimentally determined PAM NNNCC (*44*). Computational modeling shows that the fourth and fifth positions strongly prefer C’s while the second position marginally favor T.

Overall, compared with the sequence logos with 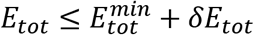 alone, the plots with 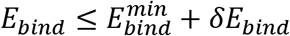 alone better recapitulated those with both 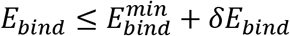 and 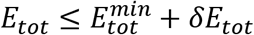 (**Fig. 4**, **Supplementary Figs. 1** and **2**), suggesting that UniDesign binding energy is a better indicator than total energy for PAM preference modeling.

In sum, the computationally predicted consensus PAM profiles by UniDesign largely recapitulated the experimentally determined consensus PAMs (**Fig. 4** and **Table 1**). Our data demonstrate that UniDesign is a useful tool to quantitatively interrogate the molecular level integrations between the native PIAAs and different PAM combos.

### Computational redesign of Cas9s reveals that PDB PAMs prefer native PIAAs

On the other hand, we want to know whether the native PIAAs are the most preferred amino acids by PDB PAMs and more generally, the consensus PAMs. We first used UniDesign to redesign the PIAAs for each Cas9 with a fixed PDB PAM. In the redesign process, the evolutionary module was disabled and only UniEF was used for energy calculation. The results are described below and summarized in **Table 2**.

**Table 2.** Summary of PIAA recapitulation results with PDB PAMs.

| Cas9 | PDB | Redesigned PIAA residue types | Recovery | Similarity |
| --- | --- | --- | --- | --- |
| SpCas9 | 4UN3 | K1107K, E1219E, R1333R, R1335R | 4/4 | 4/4 |
|  | 5F9R | K1107K, E1219E, R1333R, R1335R | 4/4 | 4/4 |
| SaCas9 | 5AXW | N985N, N986N, L989L, R991R, R1002K, R1015R | 5/6 | 6/6 |
|  | 5CZZ | N985N, N986N, L989H, R991R, R1002K, R1015R | 4/6 | 5/6 |
| FnCas9 | 5B2O | R1241R, E1449N, D1472D, S1473S, R1474R, S1555S, R1556R, R1585R, Y1586Y | 8/9 | 8/9 |
| Nme1Cas9 | 6JDV | Q981Q, H1024H, T1027S, N1029T, E1048E, G1049G | 4/6 | 5/6 |
| Nme2Cas9 | 6JE3 | K843K, D1028E, R1033R, Y1035W, K1044K | 3/5 | 5/5 |
| CdCas9 | 6JOO | I47R, K818K, F1011F, K1015Q, R1017R, R1042K, P1043E,<br>L1046L | 4/8 | 5/8 |
| St1Cas9 | 6M0W | F673Y, T1048T, M1049Q, Y1055Y, S1082S, Q1084Q, K1086Q | 4/7 | 5/7 |
| AceCas9 | 6WBR | R55R, E1044E, R1048R, R1088R, R1091R | 5/5 | 5/5 |
| Total | N/A | N/A | 37(36)/50 | 43(42)/50 |

Only one scaffold for SpCas9 and SaCas9 was counted for calculating the total recovery and similarity rates. The values outside and inside the parentheses were obtained with scaffold 5AXW or 5CZZ counted respectively.

All PIAAs for SpCas9 (4UN3 and 5F9R) and AceCas9 (6WBR) were successfully recovered. UniDesign confirmed that the native PIAAs in both Cas9s led to the lowest energy scores.

For SaCas9, four PIAAs, i.e., Asn985, Asn986, Arg991, and Arg1015 were chosen as native types, while Arg1002 was designed into a physiochemically similar Lys on both scaffolds (5AXW and 5CZZ). An extra mutation, i.e., Leu989→His, was chosen on 5CZZ.

For FnCas9 (5B2O), eight out of nine native PIAAs were picked by UniDesign, with the exception of residue Glu1449. UniDesign suggested Asn at this position.

For Nme1Cas9 (6JDV), four out of the six native PIAAs were recapitulated by UniDesign. At the other two PIAAs (Thr1027 and Asn1029), UniDesign suggested Ser and Thr, respectively, both of which do not change amino-acid polarity.

Of the five native PIAAs in Nme2Cas9 (6JE3), three were recapitulated by UniDesign; the two differences were at D1028→E and Y1035→W, where UniDesign suggested residues with similar properties.

For CdCas9 (6JOO, four of the eight native PIAAs were suggested by UniDesign. For the other four PIAAs, UniDesign suggested Lys instead of the native Arg at position 1042, Gln instead of the native Lys at position 1015, and Arg instead of the native Ile at position 47. The UniDesign suggested Arg47 is located at a solvent-exposed position that can form π-π stacking interaction with PAM nucleotide G1.

Lastly, for St1Cas9 (6M0W), four out of seven native PIAAs, i.e., Thr1048, Tyr1055, Ser1082, Gln1084, and one physiochemically similar mutation F673→Y were suggested by UniDesign. Computational modeling suggested that the mutation M1049→Q formed a 2.9 Å hydrogen bond with the PAM’s sixth A, and K1086→Q formed two hydrogen bonds (3.0 and 3.2 Å respectively) with the fourth G of PAM, while in the experimental structure, the native Lys1086 also forms two hydrogen bonds (2.8 and 2.4 Å respectively) with the PAM’s fourth G.

Overall, greater than 70% native PIAAs from all eight Cas9s were recapitulated by UniDesign. Considering the amino acids similarity (e.g., R↔K, F↔Y↔W, D↔E, and S↔T), greater than 80% PIAAs could be regarded as recovered (see **Table 2**). Such high recapitulation rates indicate that PDB PAMs show a high preference for native or native-like PIAAs, and that UniDesign is a reliable tool to predict PIAAs given a fixed PAM.

### UniDesign analysis suggests PIAA changes to accommodate different PAM sequences

For practical reasons, not all the PAMs will be used in structural biology experiments to determine their integrations with the PIAAs. In this regard, a CPD method, such as UniDesign, serves as an alternative to the structural biology experiments to scan all consensus PAMs and provide computational insights into their interactions with PIAAs. In particular, we ask the question whether for each different PAM (e.g., AGG *versus* CGG), a different combo of PIAAs will be required for favorable Cas9–DNA binding to achieve high gene-editing efficiency.

Here for each Cas9 structure, the models bearing all possible consensus PAM combos were used as a scaffold to interrogate the effects of mutations of the PIAAs on the energy scores in UniDesign. The results from different combos were combined for sequence logo analysis. For instance, four PAM combos of SpCas9 (PAM: NGG) with AGG, CGG, GGG, or TGG PAMs were used for PIAA redesign individually. Similarly, 64 (= 4 × 4 × 2 × 2) PAM combos for SaCas9 (PAM: NNGRRT) were fed into UniDesign to redesign the corresponding PIAAs.

The UniDesign results were then used to generate sequence logo plots. It is shown that in general, for most Cas9 proteins, especially SpCas9, SaCas9, FnCas9, and Nme1Cas9, all or a very high ratio of native PIAAs were computationally recapitulated based on the UniDesign computed energy scores, indicating that native PIAAs are sufficient for all consensus PAM variants (**Fig. 5**). Compared with the PIAAs recapitulation results with PDB PAMs (**Table 2**), identical design results were obtained for SpCas9 (**Fig. 5 a** and **b**), SaCas9 (**Fig. 5c** and **d**), and Nme1Cas9 (**Fig. 5f**) with non-PDB, consensus PAMs. It should also be noted that in some cases, e.g., for Nme2Cas9 (**Fig. 5g**), CdCas9 (**Fig. 5h**), and AceCas9 (**Fig. 5j**), UniDesign suggested specific PIAAs changes for different PAM variations although native or native-like amino acids were still dominating at most positions. This means that native PIAAs may not best fit some PAMs even though they are still consensus PAMs.

**Fig. 5.**
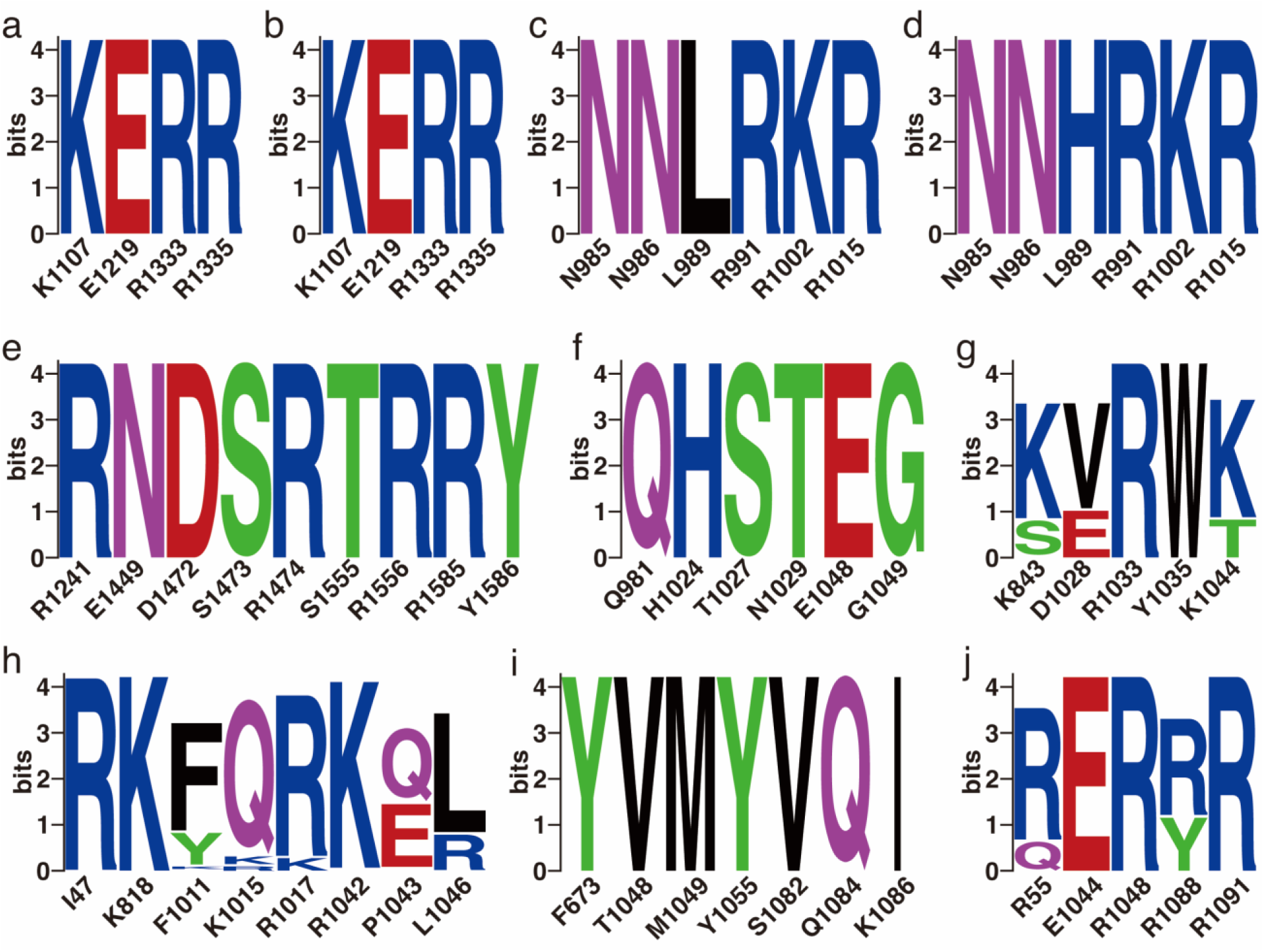
Computational redesign of Cas9 proteins reveals that consensus PAMs prefer native PAM-interacting amino acids. Illustrated are sequence logo plots for results on (**a**) scaffold 4UN3 (SpCas9) with 4 consensus PAM combos; (**b**) 5F9R (SpCas9) with 4 consensus PAM combos; (**c**) 5AXW (SaCas9) with 64 consensus PAM combos; (**d**) 5CZZ (SaCas9) with 64 consensus PAM combos; (**e**) 5B2O (FnCas9) with 4 consensus PAM combos; (**f**) 6JDV (Nme1Cas9) with 16 consensus PAM combos; (**g**) 6JE3 (Nme2Cas9) with 256 consensus PAM combos; (**h**) 6JOO (CdCas9) with 432 consensus PAM combos; (**i**) 6M0W (St1Cas9) with 32 consensus PAM combos, and (**j**) 6WBR (AceCas9) with 64 consensus PAM combos. Note that 16 and 432 consensus PAM combos were obtained for Nme1Cas9 and CdCas9 because their first two or one PAM positions were not varied respectively.

Collectively, these results indicate that, although non-PDB, consensus PAMs still highly prefer native or native-like PIAAs, a single sequence of PIAAs may not tolerate all consensus PAM combos equally, thus suggesting mutation studies on PIAAs to accommodate different PAM variations. UniDesign can serve as a useful computational tool for this purpose.

## Discussion

Understanding the PAM recognition process at the structural biology level is critical to engineering CRISPR–Cas proteins toward relaxed or altered PAM requirements, which have direct impacts on their applications in biotechnology and medicine. To this end, current studies are typically conducted as follows: first, determining the consensus PAMs; second, solving the Cas/gRNA/DNA complex structure(s) to elucidate Cas–PAM interactions; and third, structure-guided engineering of PAM requirement. In the second stage, the DNA substrate usually contained the most preferred PAM (e.g., PDB PAM as defined in this study). Despite these useful and insightful studies, a comprehensive and quantitative understanding of the relationship between a Cas and its consensus PAMs is still missing, hindering the systematic computational design of Cas9 variants with modified PAM requirements. To our knowledge, COMET is the only computational workflow designed for Cas9 PAM engineering to date (*19*). But this method is not widely used since it is inefficient and resource-costing.

In this study, we aim to develop a universal, easy-to-use CPD approach to model PNI accurately and efficiently. To test UniDesign’s effectiveness, we used it to model PAM recognition for eight Cas9 proteins. We noted that, given native PIAAs, the whole Cas9/gRNA/DNA system with a consensus PAM in general had relatively lower predicted UniDesign binding energy (and total energy) (**Fig. 3**), suggesting that PIAAs have a preference for consensus PAMs. This is consistent with experiments that Cas9 initiates DNA interrogation through specific recognition of preferred PAMs (*49, 50*). By setting *δE_bind_* and *δE_tot_* thresholds appropriately, we found that the predicted low-energy PAM combos to a large extent recapitulated the preferred consensus PAM profiles as determined by experiments (**Fig. 4**). Also, we showed that UniDesign binding energy (together with *δE_bind_*) was a better descriptor for PAM preference than total energy (together with *δE_tot_*) (**Fig. 4**, **Supplementary Figs. 1** and **2**). Conversely, we noted that, given PDB PAMs or consensus PAMs, computational redesign of PIAAs by UniDesign largely recapitulated the naturally occurring amino acids at these positions (**Table 2** and **Fig. 5**), suggesting that consensus PAMs also strongly favor native PIAA residues. Thus, computational modeling of the Cas9–PAM recognition by the UniDesign reveals the inherent mutual preference between consensus PAMs and native PIAAs resulted from long-time evolution. It should be noted that the UniDesign energy function (UniEF) was trained only on protein monomers and PPIs, but Cas9–PAM recognition modeling results suggested that the prototype UniDesign generally works on PNIs.

Another point we would like to discuss is how to use UniDesign to engineer Cas9 proteins with relaxed or altered PAMs. Based on the above analysis, we speculate that sufficient binding interaction between Cas9 and PAM is a prerequisite for DNA interrogation initiation. As a consequence, the UniDesign binding and total energy of Cas9/gRNA/DNA with PDB PAM and native PIAAs can be used as a reference/baseline. When PAM is altered (even though it is still a consensus PAM), native PIAAs may not fit the modified PAM best, resulting in a binding loss (e.g., reduced binding energy). Thus, the PIAA residues may need to be redesigned to accommodate the changed PAM, and meanwhile, other PAM-surrounding amino acids may also need to be redesigned to enhance nonspecific interactions between Cas9 and PAM to make up for the lost binding affinity. UniDesign can automate this design process to generate variants for further analysis. Also, it should be noted that UniDesign requires only minimal computational resources and can run efficiently on a personal computer and on different operating systems.

In sum, we report UniDesign, a universal CPD approach for PNI modeling and design. We demonstrate UniDesign’s effectiveness by applying it to decode Cas9–PAM recognition quantitatively for eight Cas9 proteins. This work represents the first systematic computational modeling on PAM recognition that can provide new insights for PAM engineering. We expect that UniDesign will serve as an important tool for Cas9 engineering in the field.

## Key Points

- We report UniDesign as a new universal computational protein design (CPD) framework for protein-nucleic acid interaction modeling and design.
- UniDesign is the first systematic CPD method for engineering CRISPR–Cas9 protein’s PAM requirements, and achieved good performance on diverse Cas9 proteins.
- UniDesign accurately modeled the mutual preference between natural PAMs and native PAM-interacting amino acids caused by long-term evolution
- UniDesign is fully open-sourced, computationally efficient and inexpensive, and can run on personal computers and different operating systems.

## Code availability

UniDesign is available at https://github.com/tommyhuangthu/UniDesign. All Perl scripts for this work are available at https://doi.org/10.5281/zenodo.7426027.

## Data availability

All computational data, including preprocessed Cas9 structure PDB files, PAM variant models, PIAAs repacked and redesigned models, RESFILEs, summary of PAM recognition and PIAA redesign results are available at https://doi.org/10.5281/zenodo.7426027.

## Acknowledgments

We thank members of the Y. Eugene Chen laboratory for their insightful discussions and suggestions to this study.

## References

1. P. Mali, K. M. Esvelt, G. M. Church, Cas9 as a versatile tool for engineering biology. Nat. Methods 10, 957–963 (2013).

2. P. D. Hsu, E. S. Lander, F. Zhang, Development and applications of CRISPR-Cas9 for genome engineering. Cell 157, 1262–1278 (2014).

3. A. C. Komor, A. H. Badran, D. R. Liu, CRISPR-Based Technologies for the Manipulation of Eukaryotic Genomes. Cell 168, 20–36 (2017).

4. X. Huang, D. Yang, J. Zhang, J. Xu, Y. E. Chen, Recent Advances in Improving Gene-Editing Specificity through CRISPR–Cas9 Nuclease Engineering. Cells 11, 2186 (2022).

5. L. Cong et al., Multiplex genome engineering using CRISPR/Cas systems. Science 339, 819–823 (2013).

6. D. Collias, C. L. Beisel, CRISPR technologies and the search for the PAM-free nuclease. Nat Commun 12, 555 (2021).

7. R. T. Walton, K. A. Christie, M. N. Whittaker, B. P. Kleinstiver, Unconstrained genome targeting with near-PAMless engineered CRISPR-Cas9 variants. Science 368, 290–296 (2020).

8. H. Nishimasu et al., Engineered CRISPR-Cas9 nuclease with expanded targeting space. Science 361, 1259–1262 (2018).

9. J. H. Hu et al., Evolved Cas9 variants with broad PAM compatibility and high DNA specificity. Nature 556, 57–63 (2018).

10. B. P. Kleinstiver et al., Broadening the targeting range of Staphylococcus aureus CRISPR-Cas9 by modifying PAM recognition. Nat. Biotechnol. 33, 1293–1298 (2015).

11. Y. Tian, X. Huang, Q. Li, Y. Zhu, Computational design of variants for cephalosporin C acylase from *Pseudomonas* strain N176 with improved stability and activity. Appl. Microbiol. Biotechnol. 101, 621–632 (2017).

12. J. He, X. Huang, J. Xue, Y. Zhu, Computational redesign of penicillin acylase for cephradine synthesis with high kinetic selectivity. Green Chem. 20, 5484–5490 (2018).

13. M. H. Hettiaratchi et al., Reengineering biocatalysts: Computational redesign of chondroitinase ABC improves efficacy and stability. Sci Adv 6, eabc6378 (2020).

14. L. Cao et al., Design of protein-binding proteins from the target structure alone. Nature 605, 551–560 (2022).

15. L. Cao et al., De novo design of picomolar SARS-CoV-2 miniprotein inhibitors. Science 370, 426–431 (2020).

16. A. Goldenzweig et al., Automated Structure- and Sequence-Based Design of Proteins for High Bacterial Expression and Stability. Mol. Cell 63, 337–346 (2016).

17. O. Khersonsky et al., Automated Design of Efficient and Functionally Diverse Enzyme Repertoires. Mol. Cell 72, 178–186 e175 (2018).

18. J. Ashworth et al., Computational redesign of endonuclease DNA binding and cleavage specificity. Nature 441, 656–659 (2006).

19. B. Luan, G. Xu, M. Feng, L. Cong, R. Zhou, Combined Computational-Experimental Approach to Explore the Molecular Mechanism of SaCas9 with a Broadened DNA Targeting Range. J. Am. Chem. Soc. 141, 6545–6552 (2019).

20. R. Pearce, X. Huang, D. Setiawan, Y. Zhang, EvoDesign: Designing protein-protein binding interactions using evolutionary interface profiles in conjunction with an optimized physical energy function. J. Mol. Biol. 431, 2467–2476 (2019).

21. X. Huang, R. Pearce, Y. Zhang, EvoEF2: accurate and fast energy function for computational protein design. Bioinformatics 36, 1135–1142 (2020).

22. D. Shultis et al., Changing the Apoptosis Pathway through Evolutionary Protein Design. J. Mol. Biol. 431, 825–841 (2019).

23. X. Huang, R. Pearce, Y. Zhang, *De novo* design of protein peptides to block association of the SARS-CoV-2 spike protein with human ACE2. Aging 12, 11263–11276 (2020).

24. X. Huang, R. Pearce, Y. Zhang, FASPR: an open-source tool for fast and accurate protein side-chain packing. Bioinformatics 36, 3758–3765 (2020).

25. X. Huang, R. Pearce, Y. Zhang, Toward the Accuracy and Speed of Protein Side-Chain Packing: A Systematic Study on Rotamer Libraries. J. Chem. Inf. Model. 60, 410–420 (2020).

26. H. M. Berman et al., The Protein Data Bank. Acta Crystallographica Section D Biological Crystallography 58, 899–907 (2002).

27. B. R. Brooks et al., CHARMM: A program for macromolecular energy, minimization, and dynamics calculations. J. Comput. Chem. 4, 187–217 (1983).

28. J. Huang, A. D. MacKerell, Jr., CHARMM36 all-atom additive protein force field: validation based on comparison to NMR data. J. Comput. Chem. 34, 2135–2145 (2013).

29. I. Fonfara et al., Phylogeny of Cas9 determines functional exchangeability of dual-RNA and Cas9 among orthologous type II CRISPR-Cas systems. Nucleic Acids Res. 42, 2577–2590 (2014).

30. E. Kim et al., In vivo genome editing with a small Cas9 orthologue derived from Campylobacter jejuni. Nat Commun 8, 14500 (2017).

31. M. Yamada et al., Crystal Structure of the Minimal Cas9 from Campylobacter jejuni Reveals the Molecular Diversity in the CRISPR-Cas9 Systems. Mol. Cell 65, 1109–1121 e1103 (2017).

32. R. Nakagawa et al., Engineered Campylobacter jejuni Cas9 variant with enhanced activity and broader targeting range. Commun Biol 5, 211 (2022).

33. C. Anders, O. Niewoehner, A. Duerst, M. Jinek, Structural basis of PAM-dependent target DNA recognition by the Cas9 endonuclease. Nature 513, 569–573 (2014).

34. F. Jiang et al., Structures of a CRISPR-Cas9 R-loop complex primed for DNA cleavage. Science 351, 867–871 (2016).

35. H. Nishimasu et al., Crystal Structure of Staphylococcus aureus Cas9. Cell 162, 1113–1126 (2015).

36. H. Hirano et al., Structure and Engineering of Francisella novicida Cas9. Cell 164, 950–961 (2016).

37. W. Sun et al., Structures of Neisseria meningitidis Cas9 Complexes in Catalytically Poised and Anti-CRISPR-Inhibited States. Mol. Cell 76, 938–952 e935 (2019).

38. N. Amrani et al., NmeCas9 is an intrinsically high-fidelity genome-editing platform. Genome Biol. 19, 214 (2018).

39. Z. Hou et al., Efficient genome engineering in human pluripotent stem cells using Cas9 from Neisseria meningitidis. Proc Natl Acad Sci U S A 110, 15644–15649 (2013).

40. C. M. Lee, T. J. Cradick, G. Bao, The Neisseria meningitidis CRISPR-Cas9 System Enables Specific Genome Editing in Mammalian Cells. Mol. Ther. 24, 645–654 (2016).

41. A. Edraki et al., A Compact, High-Accuracy Cas9 with a Dinucleotide PAM for In Vivo Genome Editing. Mol. Cell 73, 714–726 e714 (2019).

42. S. Hirano et al., Structural basis for the promiscuous PAM recognition by Corynebacterium diphtheriae Cas9. Nat Commun 10, 1968 (2019).

43. Y. Zhang et al., Catalytic-state structure and engineering of Streptococcus thermophilus Cas9. Nature Catalysis 3, 813–823 (2020).

44. A. Das et al., The molecular basis for recognition of 5’-NNNCC-3’ PAM and its methylation state by Acidothermus cellulolyticus Cas9. Nat Commun 11, 6346 (2020).

45. G. E. Crooks, G. Hon, J. M. Chandonia, S. E. Brenner, WebLogo: a sequence logo generator. Genome Res. 14, 1188–1190 (2004).

46. A. E. Friedland et al., Characterization of Staphylococcus aureus Cas9: a smaller Cas9 for all-in-one adeno-associated virus delivery and paired nickase applications. Genome Biol. 16, 257 (2015).

47. F. A. Ran et al., In vivo genome editing using Staphylococcus aureus Cas9. Nature 520, 186–191 (2015).

48. K. M. Esvelt et al., Orthogonal Cas9 proteins for RNA-guided gene regulation and editing. Nat. Methods 10, 1116–1121 (2013).

49. S. H. Sternberg, S. Redding, M. Jinek, E. C. Greene, J. A. Doudna, DNA interrogation by the CRISPR RNA-guided endonuclease Cas9. Nature 507, 62–67 (2014).

50. V. Globyte, S. H. Lee, T. Bae, J. S. Kim, C. Joo, CRISPR/Cas9 searches for a protospacer adjacent motif by lateral diffusion. EMBO J. 38, (2019).

