## Supplementary for "Decoding CRISPR–Cas9 PAM recognition with UniDesign"

### Contents

**Supplementary Table 1**. PAM-interacting amino acids (PIAAs) on eight CRISPR–Cas9 proteins.

**Supplementary Table 2**. UniDesign calculated $E_{bind}^{min}$ and $E_{tot}^{min}$ energy scores and the appropriately chosen $\delta E_{bind}$ and $\delta E_{tot}$ thresholds for ten CRISPR–Cas9 scaffolds.

**Supplementary Figure 1**. UniDesign predicted consensus PAMs with $E_{bind} \leq E_{bind}^{min} + \delta E_{bind}$.

**Supplementary Figure 2**. UniDesign predicted consensus PAMs with $E_{tot} \leq E_{tot}^{min} + \delta E_{tot}$.

**Supplementary Table 1**. PAM-interacting amino acids on eight CRISPR–Cas9 proteins.

| Cas9 | PDB | PAM-interacting amino acids |
|---|---|---|
| SpCas9 | 4UN3 | K1107, E1219, R1333, R1335 |
| | 5F9R | K1107, E1219, R1333, R1335 |
| SaCas9 | 5AXW | N985, N986, L989, R991, R1002, R1015 |
| | 5CZZ | N985, N986, L989, R991, R1002, R1015 |
| FnCas9 | 5B2O | R1241, E1449, D1472, S1473, R1474, S1555, R1556, R1585, Y1586 |
| Nme1Cas9 | 6JDV | Q981, H1024, T1027, N1029, E1048, G1049 |
| Nme2Cas9 | 6JE3 | K843, D1028, R1033, Y1035, K1044 |
| CdCas9 | 6JOO | I47, K818, F1011, K1015, R1017, R1042, P1043, L1046 |
| St1Cas9 | 6M0W | F673, T1048, M1049, Y1055, S1082, Q1084, K1086 |
| AceCas9 | 6WBR | R55, E1044, R1048, R1088, R1091 |

**Supplementary Table 2**. UniDesign calculated $E_{bind}^{min}$ and $E_{tot}^{min}$ energy scores and the appropriately chosen $\delta E_{bind}$ and $\delta E_{tot}$ thresholds for ten CRISPR–Cas9 scaffolds.

| Cas9 | PDB | $E_{bind}^{min}$ (UEU) | $E_{tot}^{min}$ (UEU) | $\delta E_{bind}$ (UEU) | $\delta E_{tot}$ (UEU) |
|---|---|---|---|---|---|
| SpCas9 | 4UN3 | -419.54 | -7011.84 | 3 | 3 |
| | 5F9R | -494.47 | -5109.58 | 2 | 3 |
| SaCas9 | 5AXW | -447.93 | -5767.12 | 2 | 15 |
| | 5CZZ | -456.22 | -5710.57 | 3 | 15 |
| FnCas9 | 5B2O | -668.71 | -8562.60 | 4 | 12 |
| Nme1Cas9 | 6JDV | -613.15 | -5679.16 | 5 | 8 |
| Nme2Cas9 | 6JE3 | -349.47 | -3204.96 | 3 | 15 |
| CdCas9 | 6JOO | -343.92 | -4314.55 | 3 | 15 |
| St1Cas9 | 6M0W | -614.68 | -5167.48 | 5 | 15 |
| AceCas9 | 6WBR | -377.70 | -4909.50 | 4 | 20 |

UEU: UniDesign energy units.

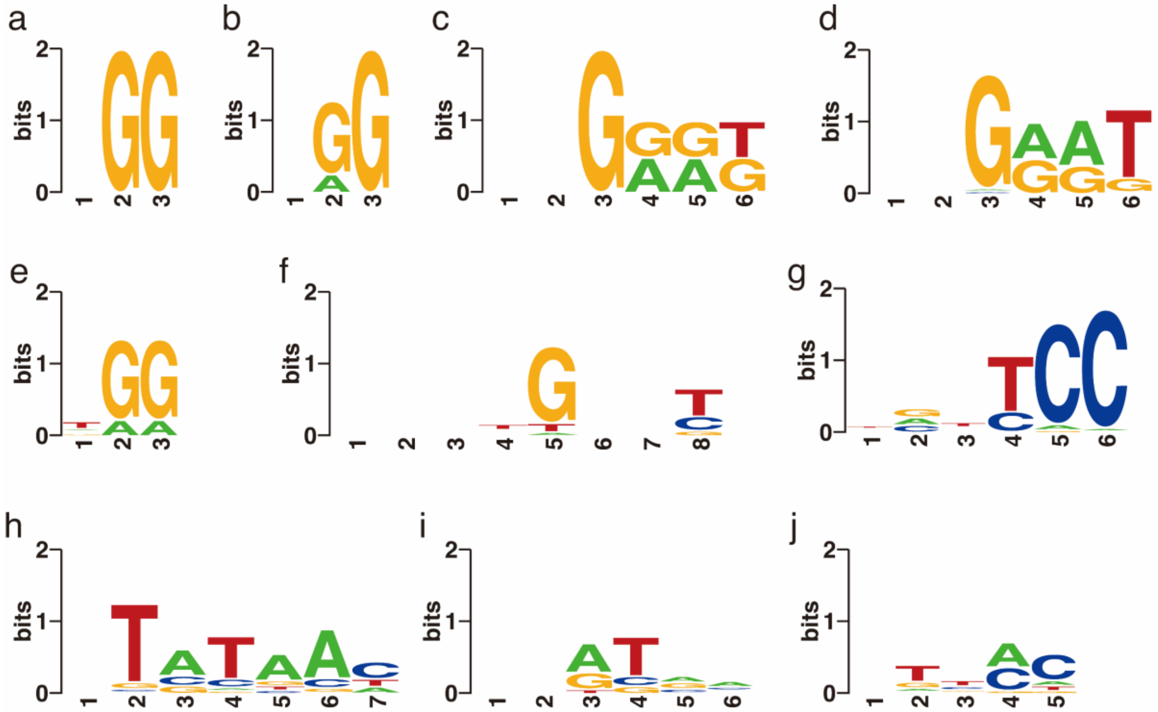


**Supplementary Figure 1**. UniDesign predicted consensus PAMs with $E_{bind} \leq E_{bind}^{min} + \delta E_{bind}$ only. The $E_{bin}^{min}$ and $\delta E_{bin}$ values are provided in Supplementary Table 2. (**a**) result on scaffold 4UN3 (SpCas9); (**b**) result on scaffold 5F9R (SpCas9); (**c**) result on scaffold 5AXW (SaCas9); (**d**) result on scaffold 5CZZ (SaCas9); (**e**) result on scaffold 5B2O (FnCas9); (**f**) result on scaffold 6JDV (Nme1Cas9); (**g**) result on scaffold 6JE3 (Nme2Cas9); (**h**) result on scaffold 6JOO (CdCas9); (**i**) result on scaffold 6M0W (St1Cas9); and (**j**) result on scaffold 6WBR (AceCas9). For Nme1Cas9 and CdCas9, the first two and one positions were excluded from PAM combo generation respectively, and the "-" symbols were manually added at these positions for plotting sequence logos.

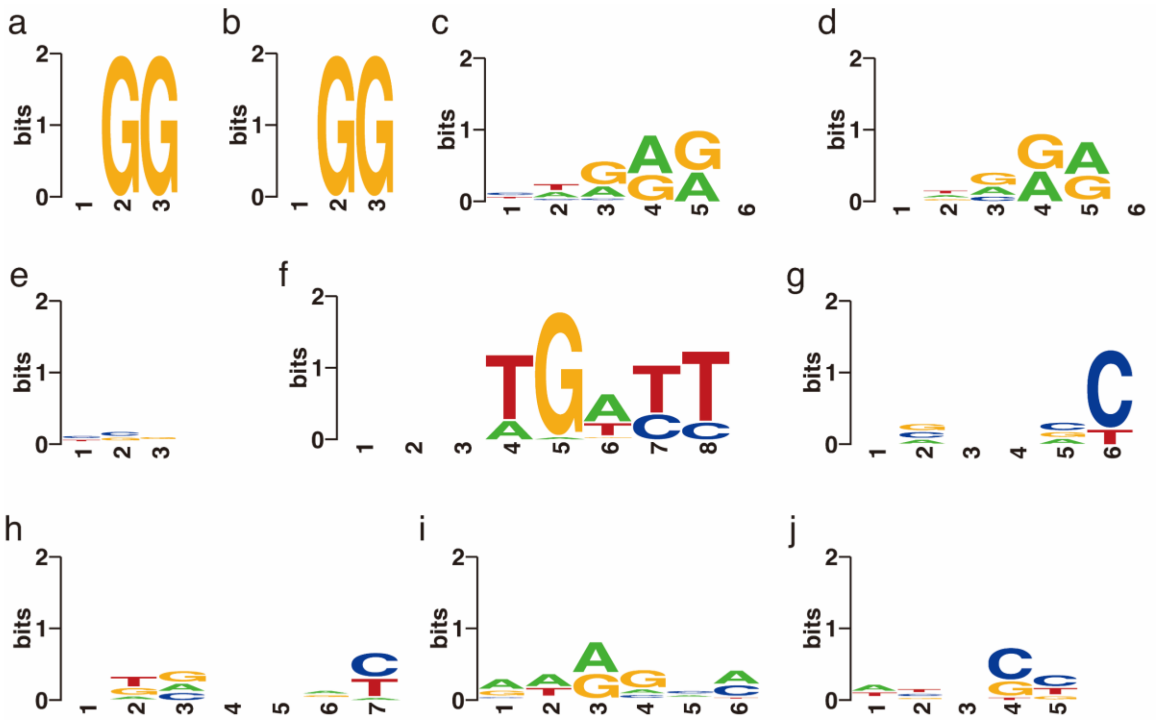


**Supplementary Figure 2**. UniDesign predicted consensus PAMs with $E_{tot} \leq E_{tot}^{min} + \delta E_{tot}$. The $E_{tot}^{min}$ and $\delta E_{tot}$ values are provided in Supplementary Table 2. (**a**) result on scaffold 4UN3 (SpCas9); (**b**) result on scaffold 5F9R (SpCas9); (**c**) result on scaffold 5AXW (SaCas9); (**d**) result on scaffold 5CZZ (SaCas9); (**e**) result on scaffold 5B2O (FnCas9); (**f**) result on scaffold 6JDV (Nme1Cas9); (**g**) result on scaffold 6JE3 (Nme2Cas9); (**h**) result on scaffold 6JOO (CdCas9); (**i**) result on scaffold 6M0W (St1Cas9); and (**j**) result on scaffold 6WBR (AceCas9). For Nme1Cas9 and CdCas9, the first two and one positions were excluded from PAM combo generation respectively, and the "-" symbols were manually added at these positions for plotting sequence logos.
